# The bone marrow niche and hematopoietic system are distinctly remodeled by CD45-targeted astatine-211 radioimmunotherapy

**DOI:** 10.1101/2025.04.04.645037

**Authors:** Matthew W. Hagen, Nicollette J. Setiawan, Shannon Dexter, Kelsey A. Woodruff, Francesca K. Gaerlan, Johnnie J. Orozco, Christina M. Termini

## Abstract

Radioimmunotherapy (RIT) is used to treat patients with hematological malignancies known to infiltrate the bone marrow (BM) microenvironment. RIT uses target-specific monoclonal antibodies stably conjugated to radionuclides to deliver cytotoxic radiation to cells of interest. While RIT is effective at delivering radiation to cancer cells, normal tissue is also exposed to radiation upon RIT, the consequences of which are largely unknown. Here, we studied the cellular and molecular effects of CD45-targeted astatine-211 (^211^At) RIT, IgG non-targeted ^211^At RIT, and Cesium-137 total-body irradiation (TBI) on hematopoietic cells and their BM niche in wild-type immunocompetent mice. Relative to non-targeted RIT or TBI, CD45-targeted RIT significantly delayed hematopoietic regeneration overall in the peripheral blood and BM and reduced hematopoietic stem/progenitor cell recovery and colony-forming ability. While BM endothelial cells (ECs) do not express the CD45 antigen, CD45-targeted RIT significantly depleted BM ECs compared to non-targeted RIT or TBI. RNA sequence analysis revealed significantly different transcriptomic profiles of BM ECs from CD45-RIT-treated mice compared to non-targeted RIT or TBI. ECs from CD45-RIT-treated mice, but not TBI or IgG-RIT-treated mice, were transcriptionally enriched for TGFβ, NOTCH, and IFNα signaling pathways compared to untreated mice. Collectively, our study indicates that CD45-targeted RIT severely impacts hematopoietic and EC niche recovery compared to non-targeted approaches. Future studies are required to determine the long-term consequences of such RIT-driven effects on BM niche physiology and how BM niche reprogramming by RIT affects cancer cells.

**KEY POINTS:**

- CD45-targeted radioimmunotherapy more effectively suppresses the hematopoietic system than non- targeted radiation delivery.
- The bone marrow vascular niche is differentially reprogrammed by CD45-targeted radioimmunotherapy compared to non-targeted radiation delivery.

## INTRODUCTION

Because of the exquisite sensitivity of hematologic malignancies to radiation, radiation therapy is pursued for some patients with blood cancers [1, 2]. Meanwhile, total body irradiation (TBI) is a longstanding myeloablative preconditioning regimen administered before hematopoietic cell transplant (HCT). Radiation causes pancytopenia, which resolves as the hematopoietic system regenerates. Ineffective hematological regeneration can leave patients more susceptible to hemorrhage or infection, which can be deadly [3, 4]. An open question in the field is how targeting radiation to distinct cellular subpopulations can mitigate or exacerbate these potentially deadly consequences.

Targeted radiation approaches, including radioimmunotherapy (RIT), can reduce off-target toxicities when treating patients with hematopoietic disorders. RIT implements radionuclides conjugated to antibodies targeting distinct antigens, such as CD45 for hematopoietic cells. While early RIT for hematologic malignancies utilized beta-emitters like iodine-131 (^131^I) or yttrium-90 (^90^Y), interest in alpha-emitters, including astatine-211 (^211^At), has grown. Unlike beta-emitters, ^211^At dispenses a high payload of radiation over a short distance (50-90µm), thereby concentrating radiation to cells of interest [5, 6]. ^211^At conjugated to anti-CD45 antibodies can be used as a nonmyeloablative conditioning mechanism to support HCT in canines [7] and humans in early-phase clinical trials [8]. Further, clinical trials using ^211^At RIT that targets CD38 and CD45 to treat patients with hematological malignancies have been undertaken [9–12]. However, the cellular and molecular consequences incurred by the non-malignant hematopoietic system upon RIT, which deposits high energy to a region the size of a few cell diameters, are poorly understood.

Here, we compare the impact of CD45-targeted and IgG non-targeted RIT on the peripheral and bone marrow (BM) hematopoietic cell compartments and how these changes compare to TBI with Cesium-137 (^137^Cs). We report distinct changes to the hematopoietic system upon CD45-targeting, including robust lymphodepletion, BM hematopoietic stem cell depletion, and vascular tissue changes. Our findings detail the profound physical, cellular, and transcriptional remodeling of the BM endothelial cell (EC) compartment upon CD45-targeting RIT that are not detected upon IgG-RIT or TBI. Our findings identify potential directions for therapeutic interventions that target the pathways dysregulated in the vascular niche to better restore this microenvironment after RIT.

## METHODS

### Study resources

Detailed information for resources used throughout this study is included in **Supplementary Table 1**.

### Radiolabeling

30F11, a rat IgG2b anti-murine CD45 antibody, was made by the Fred Hutch Antibody Production Shared Resource from hybridomas using a hollow-fiber bioreactor; isotype-matched control rat IgG2b was used (BioXCell). Antibodies were conjugated to B10-NCS [13] to facilitate stable labeling with ^211^At isolated from an irradiated bismuth target through a wet chemistry approach as described [14].

### Mouse models

Animal procedures were performed in accordance with the Fred Hutchinson Cancer Center Institutional Animal Care & Use Committee (PROTO202100049 & PROTO000001716). Mice were housed and maintained in the Fred Hutch Comparative Medicine facility. Mice were purchased from The Jackson Laboratory. *Cdh5-cre; ROSA26-tdTomato* mice were generated by crossing B6.FVB-Tg(Cdh5-cre)7Mlia/J (Strain#:006137) with B6.Cg-*Gt(ROSA)26Sor^tm14(CAG-tdTomato)Hze^*/J (Strain#:007914); F1 offspring were used in downstream applications. Female C57BL/6J (Strain#:000664) aged 9-13 weeks and mixed-sex *Cdh5-cre; ROSA26-tdTomato* aged 10-15 weeks were used. RIT mice received 100µg of anti-CD45 or IgG labeled with 20µCi of ^211^At in 200µL of PBS intravenously. Cesium irradiation was administered by single dose (500cGy) using a Mark 1 Irradiator. For kinetic studies, mice were treated in two cohorts. Mice studied at treatment day+3 are statistically grouped with and henceforth referred to as day+4; mice studied at treatment day+29 are statistically grouped with and henceforth referred to as day+28.

### Cell/tissue isolations and flow cytometry

BM was isolated from one femur and both tibias [15], lysed with ACK buffer and processed in complete IMDM (IMDM + 10%FBS + 1% penicillin-streptomycin). For hematopoietic stem/progenitor cell analyses, BM was lineage depleted using a Direct Lineage Depletion Kit. For EC analyses, femurs and tibias were isolated from mice and crushed using a mortar and pestle containing liberase [16]. Peripheral blood was collected into EDTA immediately prior to euthanasia, and complete blood counts were analyzed (Element HT5 Veterinary Hematology Analyzer). Remaining blood was lysed using ACK buffer. Lysed BM or blood was stained using antibodies or isotype controls (**Supplemental Table 1**) and analyzed by flow cytometry (LSRFortessa X-50 or FACSymphony S6). Data were analyzed using FlowJo. Cell frequencies are displayed as percent of live cells. Cell numbers are calculated using WBC (blood), whole marrow (BM), or normalized to cytometer input (ECs).

### RNA sequencing

At least 1x10^4^ ECs (CD45^-^/*Cdh5*-tdTomato^+^) were isolated using fluorescence-activated cell sorting; RNA was isolated using the RNeasy Micro Kit. RNA was sequenced by the Fred Hutch Genomics & Bioinformatics Shared Resource. cDNA was prepared using the SMART-Seq v4 Ultra Low Input RNA kit and libraries generated using NexteraXT DNA library prep kit; sequencing was performed on a NovaSeqX+ instrument (NOVASeqX+ 10B lane, 1250M reads). For analysis, STAR with 2-pass mapping was used to align paired-end reads to mm10 genome assembly. FastQC, RNA-SeQC, and RSeQC were used for QC. edgeR was used to detect differential gene expression between groups. Genes with low expression were excluded using function filterByExpr with min.count=10 and min.total.count=15. The filtered expression matrix was TMM-normalized and subjected to significance testing using edgeR’s quasi-likelihood pipeline [17]. A gene was deemed differentially expressed if it met the cutoffs of log2 fold change, and Benjamini-Hochberg adjusted p-values. Gene set enrichment analysis (GSEA) was performed by function fgsea within package fgsea [18]. Data are deposited in GEO (GSE293137).

### Colony forming unit assays

BM cells were dissolved in 600µL of complete IMDM and added to 3mL of MethoCult GF M3434; 1.2mL of this solution was added to 2 wells in SmartDishes. Cells were grown in an incubator (37°C, 5% CO2) for 14 days and counted using a STEMvision automated colony counting machine using the human 14-day algorithm with colony misidentifications corrected by a single observer. Averages of 2 wells per mouse are reported.

### Histology and microscopy

One femur per mouse was fixed in formalin for 72 hours, washed with PBS, and decalcified for 14 days with 0.5M EDTA at 4°C. Decalcified femurs were paraffin-embedded, and 4 µm sections generated at the Fred Hutch Experimental Histopathology Shared Resource. Sections were stained using hematoxylin and eosin or anti-CD31 with nuclear counterstain. Full-section scans were generated using an Aperio VERSA200 digital pathology scanner with a 20X objective. CD31 images were analyzed by creating serial random forest classifiers using HALO. Broad regions of interest encompassing the BM compartment, excluding the epiphysis, were defined by a single observer. A primary classifier was trained to identify glass, cortical bone, fold and tear artifacts, and central marrow. Central marrow was passed to a secondary classifier trained to identify CD31^+^ vascular tissue. Vascular area fraction was defined as the total CD31^+^ vascular tissue area divided by the total central marrow area in a sample. Both classifiers were trained by a single observer on samples across treatment groups and time points.

### Statistics

Hypothesis tests were performed using Prism. Non-genomic multi-timepoint data were analyzed using two-way ANOVA against treatment, timepoint, and interaction. When interactions were significant, Holm- Šídák’s multiple comparisons of CD45-RIT and other treatments were performed. Single timepoint data were analyzed with one-way ANOVA and Holm-Šídák’s multiple comparisons of all groups.

## RESULTS

### Targeted and non-targeted radiation delivery differentially deplete PB hematopoietic cells

We studied how the peripheral hematopoietic system is affected by targeted and non-targeted RIT employing the α-emitting radionuclide ^211^At conjugated to anti-CD45 or an IgG isotype control. Adult C57BL/6 mice were treated with 20µCi of IgG-RIT, CD45-RIT, or sublethal TBI (500cGy ^137^Cs) as a non-RIT radiation comparator. We isolated the peripheral blood (PB) at day +1, 4, 7, 14, or 28 post-treatment for analysis (**Figure 1A**).

**Figure 1:**
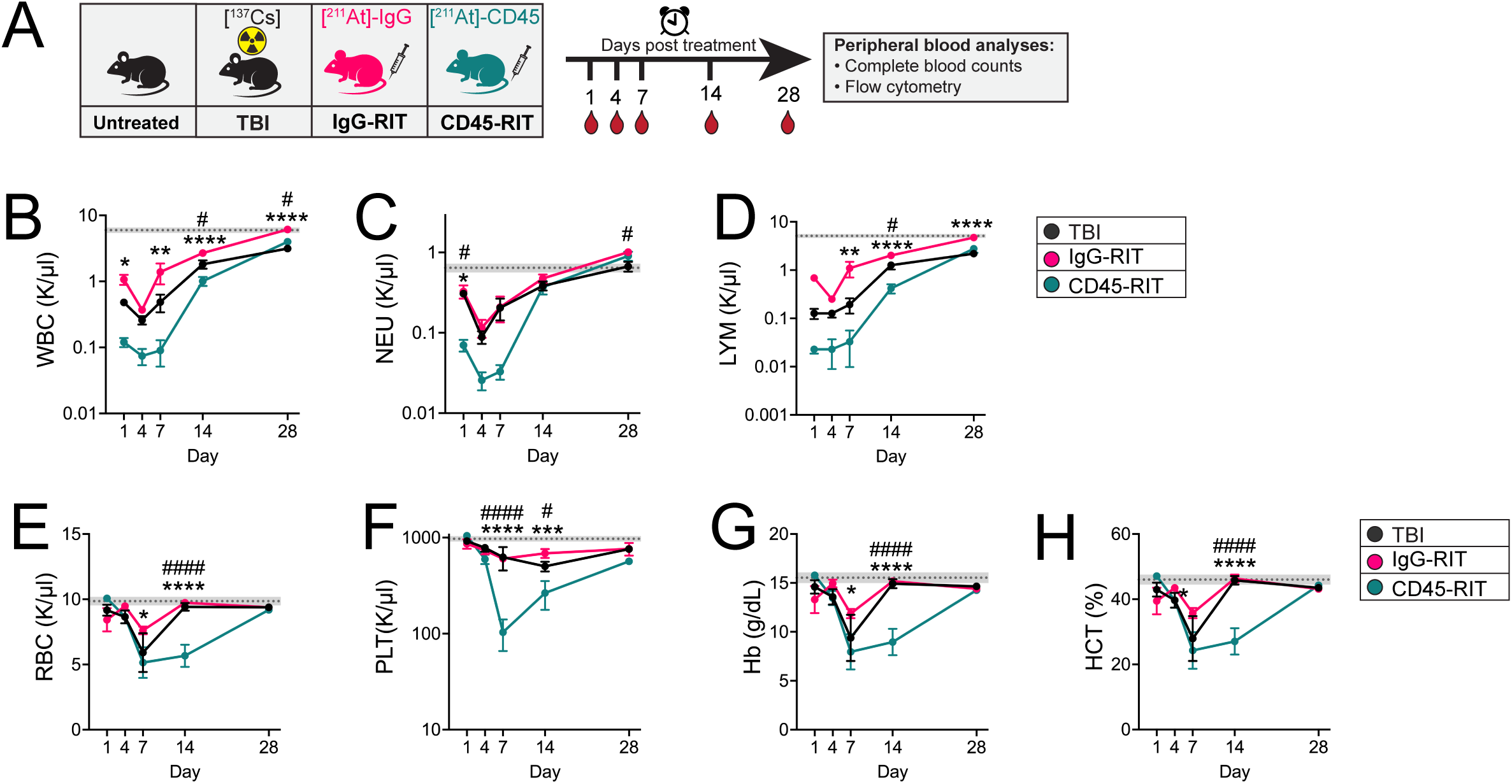
Targeted and non-targeted radiation delivery differentially deplete peripheral hematopoietic cells. **(A)** Diagram depicting the experimental design used to examine the impact of targeted or non-targeted RIT on the PB; each mouse was sampled once. Complete blood count analyses from PB for **(B)** white blood cells, **(C)** neutrophils, **(D)** lymphocytes, **(E)** red blood cells, **(F)** platelets, **(G)** hemoglobin, and **(H)** hematocrit in untreated, TBI-treated, IgG-RIT-treated, and CD45-RIT-treated mice. (n=7-9 mice per condition per time point from 2 experimental cohorts; dotted line and shaded bar indicate mean ± SEM untreated levels; statistics denote two-way ANOVA test followed by Holm-Šídák’s multiple comparisons test; \**P* < .05; \*\**P* < .01; \*\*\**P* < .001; \*\*\*\**P<*.0001 for CD45-RIT vs. IgG-RIT; #*P* < .05, ###*P<*.001 for CD45-RIT vs. TBI).

There were significantly fewer white blood cells at day +1 after CD45-RIT compared to IgG-RIT. All groups reached nadir at day +4 after treatment, and white blood cellularity remained significantly lower in CD45-RIT- treated mice compared to IgG-RIT through day +28 (**Figure 1B**). Neutrophils were significantly depleted at day +1 after CD45-RIT treatment compared to IgG-RIT or TBI groups (**Figure 1C**). PB lymphocytes were significantly reduced in CD45-RIT-treated mice compared to IgG-RIT-treated mice at days +7-28 after treatment (**Figure 1D**). At day +14, CD45-RIT-treated mice had fewer lymphocytes than both IgG-RIT and TBI groups. These data suggest that CD45-targeted RIT delays white blood cell and lymphocyte recovery after radiation exposure compared to non-targeted approaches.

PB red blood cells, platelets, hemoglobin, and hematocrit levels were also significantly decreased at day +14 after CD45-RIT treatment compared to IgG-RIT or TBI, but these differences resolved by day +28 (**Figure 1E-H**). There was a pronounced reduction in platelet count at day +7 and +14 after CD45-RIT treatment compared to IgG-RIT or TBI (**Figure 1H**), suggesting CD45 antigen specificity promoted thrombocytopenia compared to non-targeted radiation. These data indicate that CD45-RIT slows peripheral hematologic recovery compared to TBI or non-targeted IgG-RIT.

### CD45-RIT delays PB hematopoietic lineage cell recovery compared to non-targeted RIT

We next examined whether defects in PB recovery upon CD45-targeting RIT were confined to a specific hematopoietic lineage using flow cytometry (**Figure 2A-D**). PB B220^+^ B cell and CD3^+^ T cell frequencies were significantly decreased upon CD45-RIT at day +1 and +7 after treatment compared to IgG-RIT (**Figure 2E-F**), concomitant with elevated Mac-1/Gr-1^+^ myeloid cell frequency (**Figure 2G**). PB Ter-119^+^ erythroid cell frequencies were decreased upon CD45-RIT compared to TBI between days +14 and +28 (**Figure 2H**). These findings highlight the unique impacts of CD45-targeted RIT on the PB hematopoietic cell compartment.

**Figure 2:**
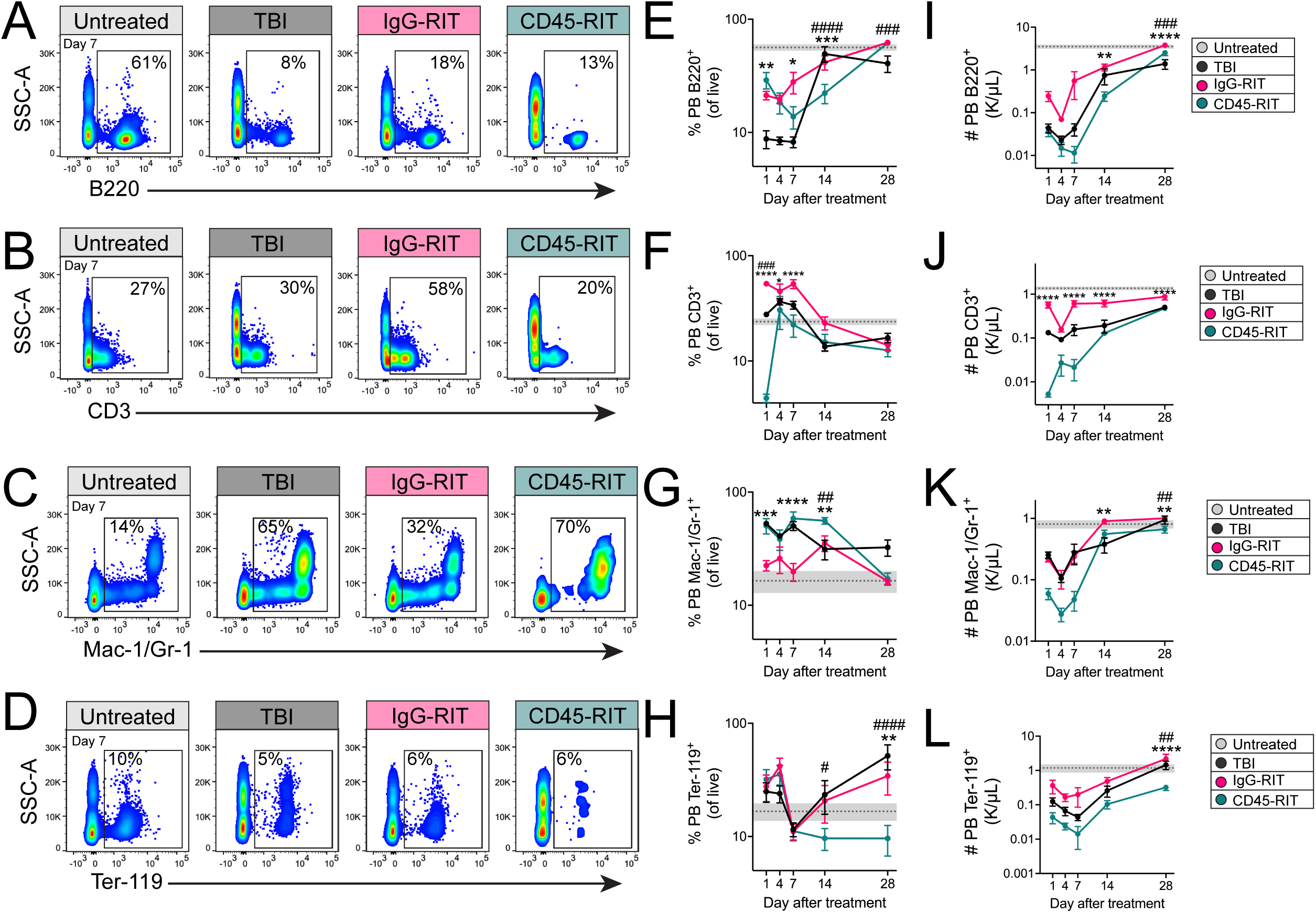
CD45-RIT delays peripheral hematopoietic lineage cell recovery compared to non-targeted RIT. Representative flow cytometry scatter plots depicting the gating strategy to detect **(A)** B220^+^ B cells, **(B)** CD3^+^ T cells, **(C)** Mac-1/Gr-1^+^ myeloid cells, and **(D)** Ter-119^+^ erythroid cells in the PB of mice after TBI, IgG-RIT, or CD45-RIT treatment at treatment day +7. The frequency of **(E)** B220^+^ B cells, **(F)** CD3^+^ T cells, **(G)** Mac-1/Gr-1^+^ myeloid cells, and **(H)** Ter-119^+^ erythroid cells was calculated. The numbers of peripheral blood **(I)** B220^+^ B cells, **(J)** CD3^+^ T cells, **(K)** Mac-1/Gr-1^+^ myeloid cells, and **(L)** Ter-119^+^ erythroid cells were quantified by multiplying relative frequencies by white blood cell counts. (n=7-9 mice per condition per time point from 2 experimental cohorts; dotted line and shaded bar indicates mean ± SEM untreated levels; statistics denote two-way ANOVA test followed by Holm-Šídák’s multiple comparisons tests; \**P* < .05; \*\**P* < .01; \*\*\**P* < .001; \*\*\*\**P<*.0001 for CD45- RIT vs. IgG-RIT; #*P* < .05, ###*P<*.001, ####*P<*.0001 for CD45-RIT vs. TBI)

PB hematopoietic cell numbers also varied by treatment type. At day +14 after CD45-RIT treatment, there was a significant reduction in B220^+^ B cells, CD3^+^ T cells, and Mac-1^+^/Gr-1^+^ myeloid cell numbers compared to IgG- RIT samples, while Ter-119^+^ erythroid cell numbers were not different (**Figure 2I-L**). Notably, PB CD3^+^ T cells were significantly lower upon CD45-RIT treatment compared to IgG-RIT as early as day +1 after treatment, and this reduction was evident through day +28. Therefore, compared to non-targeted RIT, CD45-targeted RIT more robustly depletes PB B, T, and myeloid cells.

### CD45-targeted RIT delays BM regeneration

Because the BM is the primary site of hematopoiesis in adults, we investigated how CD45-targeted RIT affects BM recovery (**Figure 3A**). At day +1 after treatment, CD45-RIT-treated mice had significantly fewer BM cells than IgG-RIT-treated mice (**Figure 3B**). However, CD45-RIT-treated mice had more than tenfold fewer BM cells at day +7 after treatment than IgG-RIT or TBI. The significantly lower BM cellularity in CD45-RIT-treated samples compared to IgG-RIT or TBI persisted through day +28 after treatment (**Figure 3B**). These data were corroborated with histological analyses of hematoxylin and eosin-stained femurs (**Figure 3C**). These data underscore the major cellular and tissue consequences of radiation on the BM, which are far more pronounced with CD45 targeting.

**Figure 3:**
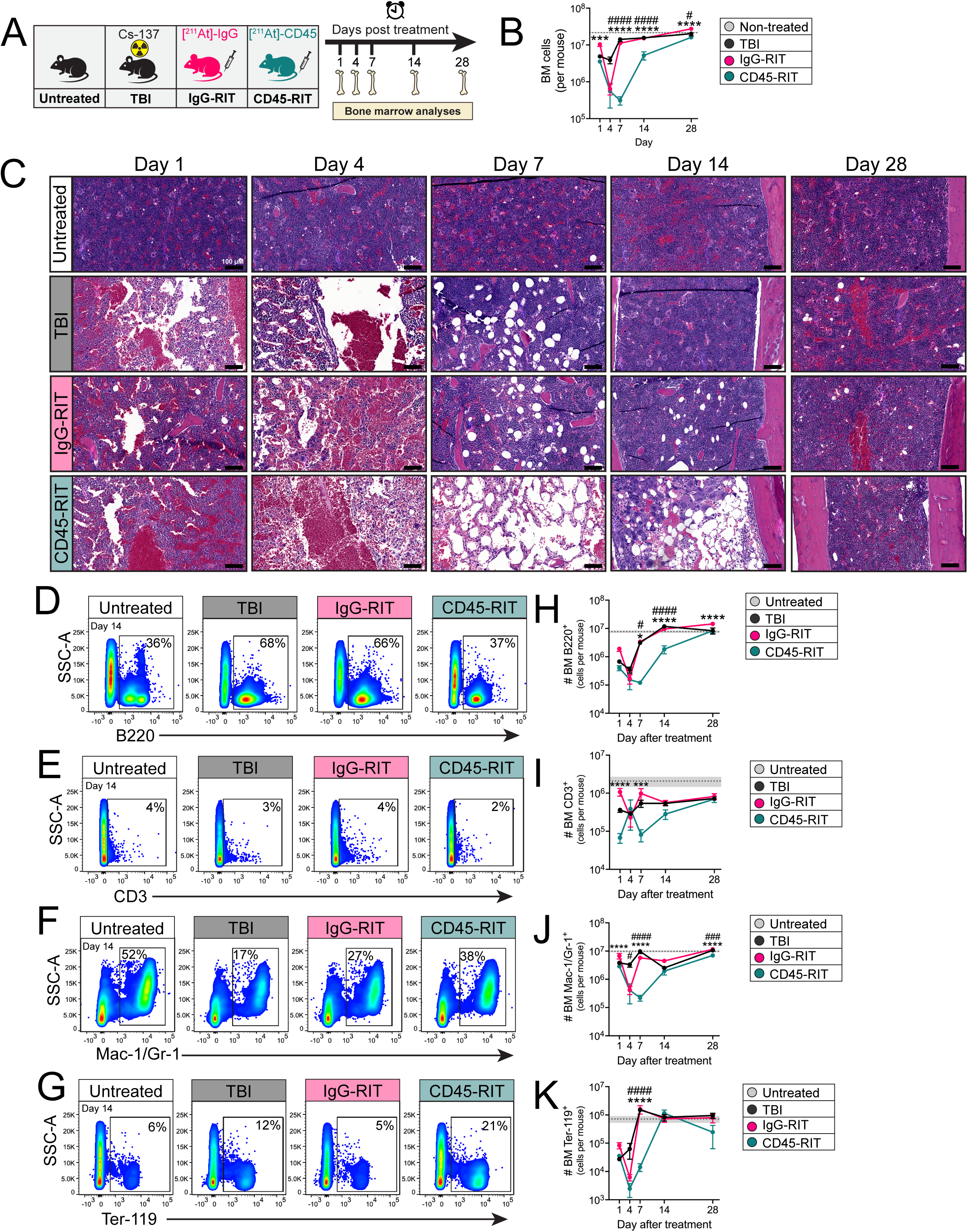
CD45-targeted RIT delays the regeneration of mature hematopoietic cells in the bone marrow. **(A)** Diagram depicting experimental design used to examine the impact of targeted or non-targeted RIT on the BM. **(B)** BM cellularity in untreated, TBI-treated, IgG-RIT-treated, and CD45-RIT-treated mice. **(C)** Hematoxylin and eosin staining of femur sections imaged using a brightfield microscope at 20X magnification are depicted. Scale bar = 100µm. Representative flow cytometry scatter plots from treatment day +14 depicting the gating strategy of **(D)** B220+ B cells, **(E)** CD3+ T cells, **(F)** Mac-1/Gr-1+ myeloid cells, and **(G)** Ter-119+ erythroid cells in the BM of mice after TBI, IgG-RIT, or CD45-RIT at day +1, +4, +7, +14, or +28 after treatment. The numbers of BM **(H)** B220^+^ B cells, **(I)** CD3^+^ T cells, **(J)** Mac-1/Gr-1^+^ myeloid cells, and **(K)** Ter-119^+^ erythroid cells were quantified by multiplying relative frequencies by BM cell counts in one femur and two tibias. (n=7-9 mice per condition per timepoint from 2 experimental cohorts; dotted line and shaded bar indicates mean ± SEM untreated levels; statistics denote two-way ANOVA test followed by Holm-Šídák’s multiple comparisons test of CD45-RIT compared to IgG-RIT or TBI; \**P* < .05; \*\*\**P* < .001, \*\*\*\**P<*.0001 for CD45-RIT vs. IgG-RIT; #*P* < .05, ###*P<*.001, ####*P<*.0001 for CD45-RIT vs. TBI).

### CD45-targeted RIT more severely depletes BM hematopoietic cells than non-targeted approaches

Flow cytometry was used to quantify BM B220^+^ B cells, CD3^+^ T cells, Mac-1/Gr-1^+^ myeloid cells, and Ter-119^+^ erythroid cells upon treatment (**Figure 3D-G**). BM B220^+^ B cells were significantly reduced in CD45-RIT-treated samples compared to IgG-RIT or TBI at day +7 and +14 after treatment (**Figure 3H**), while CD3^+^ T cells were reduced at day +1 and +7 compared to IgG-RIT (**Figure 3I**). Significantly lower BM myeloid and erythroid cells were apparent in CD45-RIT-treated samples compared to IgG-RIT or TBI at day +7 after treatment (**Figure 3J-K**). BM erythroid cell differences between groups resolved by day +14 after treatment, while myeloid cells remained modestly reduced at day +28 after CD45-RIT compared to IgG-RIT, suggesting delayed recovery of myeloid cell homeostasis upon targeted RIT. These data highlight that CD45-targeted RIT causes more severe BM hematopoietic cell depletion than non-targeted approaches.

### BM hematopoietic stem and progenitor cells are depleted by CD45-targeted RIT

Hematopoietic stem cells (HSCs) also occupy the BM niche, where they give rise to progenitors and hematopoietic lineage cells. Previous work suggests that HSCs are more resistant to apoptosis upon TBI than more proliferative hematopoietic cells [19, 20]. However, the radiosensitivity of HSCs to RIT is unknown. We, therefore quantified the BM c-Kit^+^Sca-1^+^Lineage^-^ (KSL) hematopoietic stem/progenitor cells (HSPCs) after TBI, IgG-RIT, or CD45-RIT treatment (**Figure 4A**). As early as day +1 after treatment, CD45-RIT-treated BM had significantly fewer KSL cells than IgG-RIT treated (**Figure 4B**). At day +14 after treatment, CD45-RIT-treated BM had significantly fewer KSL cells compared to TBI or IgG-RIT treated. We analyzed a further purified population of KSL cells negative for CD34 (34^-^KSL cells), which enriches for long-term HSCs [21]. CD45-RIT- treated mice had nearly tenfold fewer BM 34^-^KSL HSCs at day +14 after treatment than IgG-RIT and TBI (**Figure 4C**). These data highlight the robust capacity for CD45-RIT to deplete BM HSPCs compared to non-targeted approaches.

**Figure 4:**
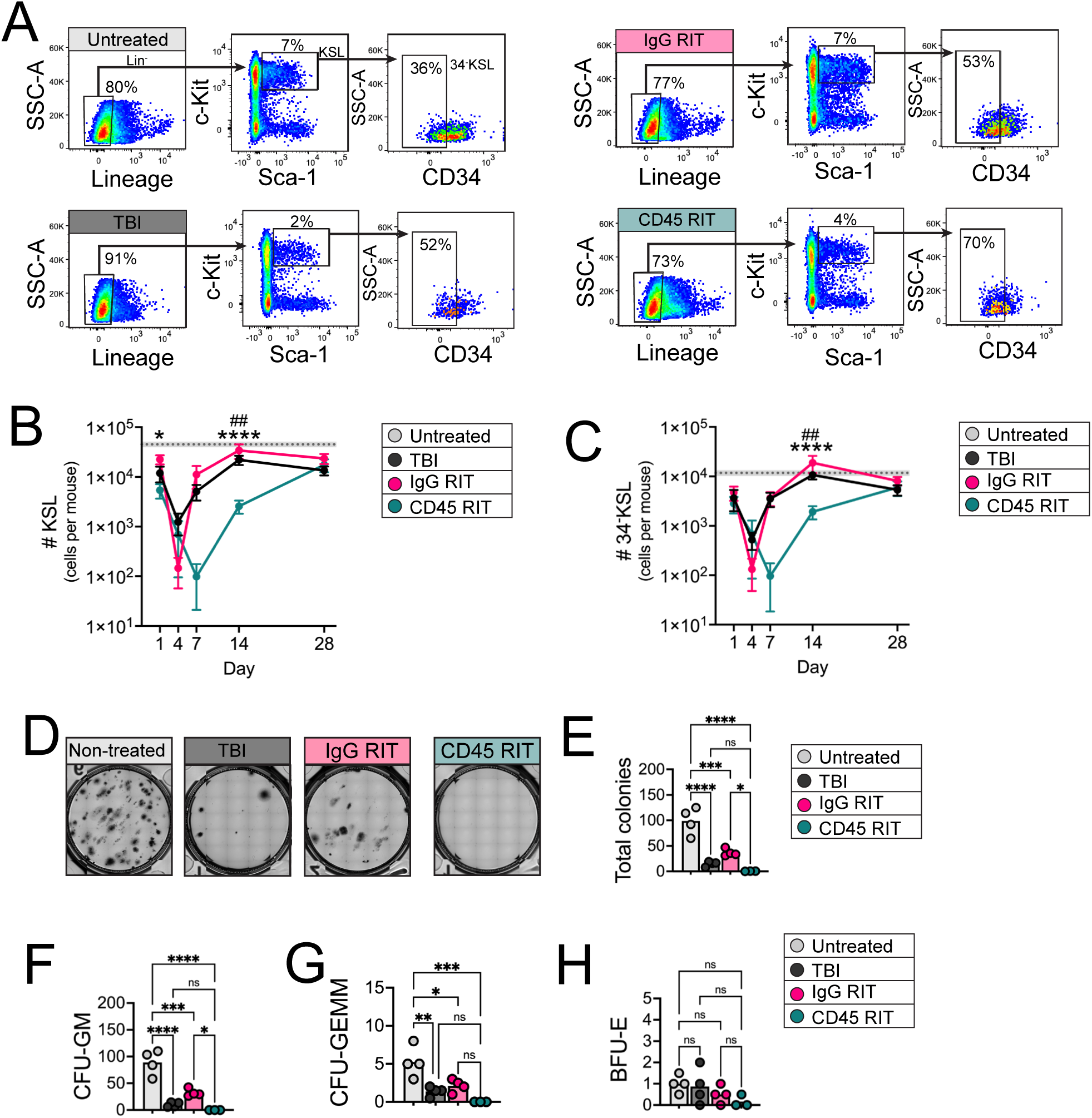
Bone marrow hematopoietic stem and progenitor cells are depleted by CD45-targeted RIT. **(A)** Representative scatter plots depicting flow cytometric gating strategy of BM HSPCs in untreated, TBI -treated, IgG-RIT-treated, and CD45-RIT-treated mice at day +14 after TBI, IgG-RIT, or CD45-RIT treatment or untreated mice. **(B)** The numbers of KSL cells per mouse was quantified by multiplying the KSL cell frequency overall by bone marrow cellularity in one femur and two tibias. **(C)** The numbers of 34^-^KSL cells were quantified by multiplying 34^-^KSL cell frequency overall by bone marrow cellularity in one femur and two tibias. **(D)** Images depicting colony forming cell potential after +14 days of culture of whole bone marrow cells isolated from treated or untreated mice at day +8 after treatment and quantification of **(E)** total colonies, **(F)** CFU-GM colonies, **(G)** CFU-GEMM colonies, and **(H)** BFU-E colonies. (For HSPC flow cytometry, n=7-9 mice per condition per timepoint from 2 experimental cohorts; dotted line and shaded bar indicates mean ± SEM untreated levels; statistics denote two-way ANOVA test followed by Holm-Šídák’s multiple comparisons test of CD45-RIT compared to IgG-RIT or TBI; \**P* < .05; \*\*\*\**P<*.0001 for CD45-RIT vs. IgG-RIT; ##*P<*.01 for CD45-RIT vs. TBI; for colony forming cell analyses, statistics denote one-way ANOVA followed by predefined post-hoc Holm-Šídák’s comparisons of untreated versus each treatment and IgG versus CD45-RIT from n=3-4 mice per condition; ns, not significant, \**P* < .05; \*\**P* < .01; \*\*\**P* < .001; \*\*\*\**P<*.0001. Displayed values are averages of 2 technical replicates per mouse).

A defining function of HSPCs is their differentiation ability, which we measured by testing the colony-forming unit (CFU) potential of BM cells from treated mice. CD45-RIT-treated BM had significantly reduced colony-forming potential compared to untreated and IgG-RIT-treated BM (**Figure 4D-E**). There was impaired CFU granulocyte-macrophage (CFU-GM) and granulocyte, erythrocyte, monocyte, megakaryocyte (CFU-GEMM) potential upon CD45-RIT treatment compared to IgG-RIT (**Figure 4F-G**). Blast-forming unit erythrocyte (BFU-E) colonies were low in frequency and not significantly different between treatments (**Figure 4H**). Thus, CD45-targeted RIT impairs BM colony-forming ability compared to IgG-RIT.

### The BM vascular niche is remodeled by CD45-targeted RIT

The BM endothelial niche is critical for supporting hematopoiesis during homeostasis but becomes significantly remodeled during hematologic injuries incurred during chemotherapy or irradiation [22–26]. We next sought to determine whether ^211^At administration via targeted or non-targeted methods similarly affects the BM EC niche. We performed immunohistochemical analyses for the EC antigen, CD31, on femur sections from untreated mice or mice treated with TBI, IgG-RIT, or CD45-RIT.

CD31 immunohistochemical staining showed major restructuring of the vascular niche upon all treatments compared to untreated marrow, which retained sinusoidal and arteriolar vascular structures that are thin and contiguous (**Figure 5A**). At day +1 after treatment, the vessels of treated mice were noticeably enlarged compared to untreated controls, and this intensified at day +4. Vascular enlargement appeared to resolve by day +7 in both non-targeted groups but remained severe in the CD45-RIT group. All groups showed resolution by day +14 (**Figure 5A**). Quantitative analysis reflects this: the BM vascular fraction was significantly higher in the CD45 group than TBI at day +4 and higher than either non-targeted group at day +7 (**Figure 5B**). These data indicate that the BM vascular niche is differentially remodeled by targeted and non-targeted RIT.

**Figure 5:**
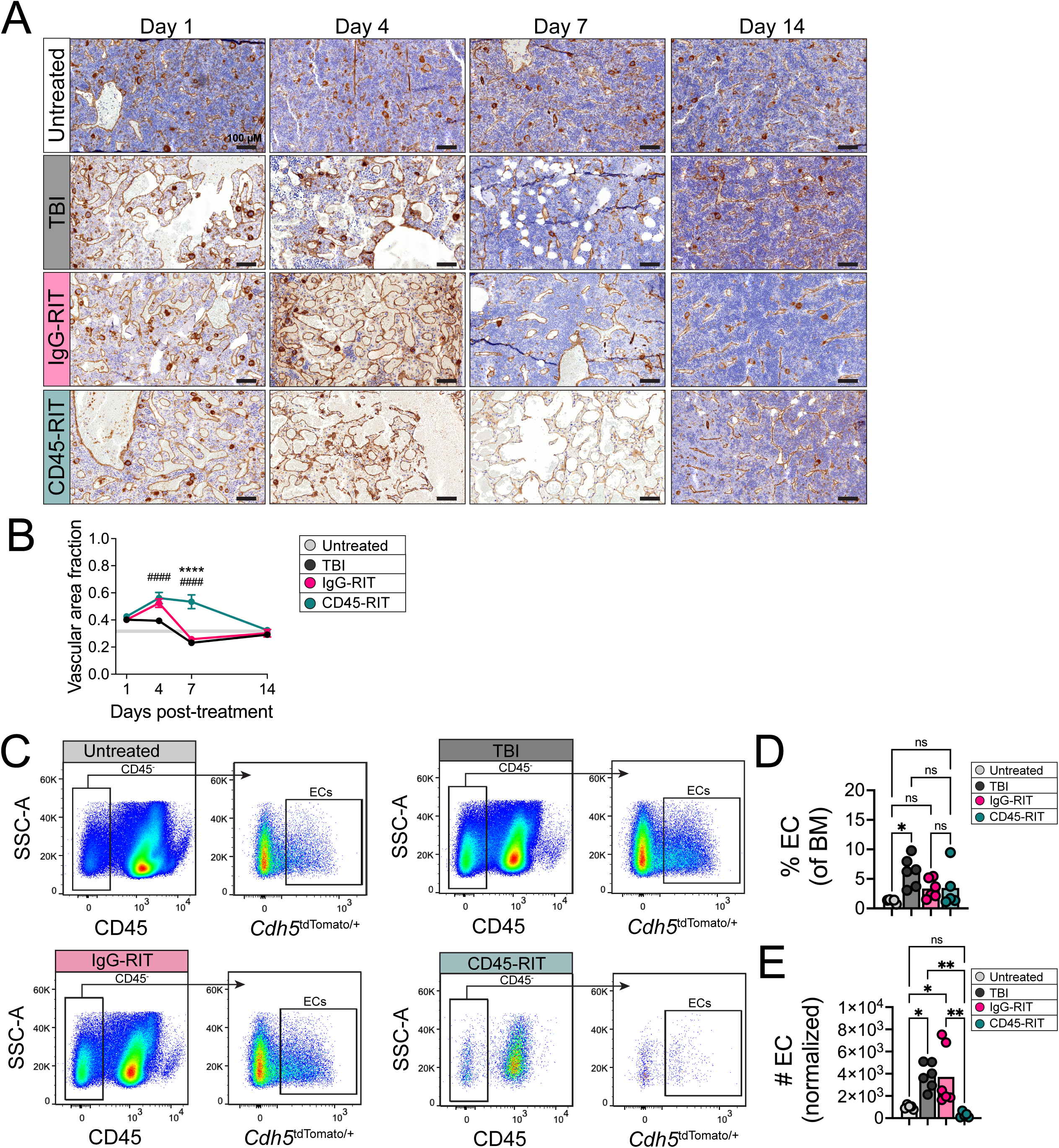
The vascular bone marrow niche is remodeled by CD45-targeted RIT. **(A)** Immunohistochemical analysis of femur sections for CD31 staining in untreated, TBI, IgG-RIT-treated, and CD45-RIT-treated mice at day +1, +4, +7 and +14 after treatment; scale bars 100µm. **(B)** The vascular area fraction was quantified using HALO software at each timepoint analyzed (n=4-9 mice per condition per timepoint from n=2 independent experiments; statistics denote two-way ANOVA test followed by Holm-Šídák’s multiple comparisons test of CD45-RIT compared to IgG-RIT or TBI; \*\*\*\**P*<.0001 for CD45-RIT vs. IgG-RIT; ####*P*<.0001 for CD45-RIT vs. TBI). **(C)** Representative flow cytometry gating of CD45^-^tdTomato^+^ cells from *Cdh5-cre; ROSA26-tdTomato* mice treated with TBI, IgG-RIT, or CD45-RIT and analyzed at day +8 after treatment. **(D)** The frequency of CD45^-^ tdTomato^+^ cells was quantified at day +8 after treatment. **(E)** CD45^-^tdTomato^+^ cell numbers per sample were quantified as the number of EC events per 200,000 events run on the cytometer. (n=5-6 mice per condition per timepoint from n=3 independent experiments; statistics denote one-way ANOVA and Holm-Šídák’s multiple comparisons tests; ns, not significant, \**P* < .05; \*\**P* < .01; \*\*\*\**P<*.0001).

### CD45-targeted RIT depletes BM ECs

To build upon the visual evidence that the BM vasculature is differentially remodeled by CD45-targeted RIT, flow cytometry was used to quantify BM EC abundance +8 days after treatment. As previously reported, TBI with ^137^Cs significantly increased EC frequency compared to untreated mice (**Figure 5C-D**) [25]. However, IgG-RIT and CD45-RIT did not increase EC frequency compared to untreated mice (**Figure 5C-D**). EC abundance was also elevated upon TBI or IgG-RIT treatment compared to untreated mice, while EC numbers upon CD45-RIT treatment were unchanged (**Figure 5E**). These data suggest that CD45-targeting differentially affects the abundance of BM ECs compared to TBI or non-targeted IgG-RIT approaches.

### The BM vascular niche is transcriptionally reprogrammed by CD45-targeted RIT

Previous reports indicate that TBI significantly remodels BM ECs not just at the tissue scale but also transcriptionally [27, 28]. BM ECs from untreated, TBI-, IgG-RIT-, or CD45-RIT-treated mice were isolated at day +8 after treatment and analyzed using bulk RNA sequencing (**Figure 6A**). Samples distinctly clustered based on principal component analysis, and CD45-RIT-treated samples cluster most distinctly from other treatments (**Figure 6B**). Compared to ECs from untreated animals, thousands of genes were differentially expressed in ECs from TBI, IgG-RIT, and CD45-RIT-treated mice (**Figure 6C-E**). ECs from CD45-RIT-treated mice also exhibited more than 2,000 differentially expressed genes compared to IgG-RIT treatment (**Figure 6F**). Therefore, targeted and non-targeted radiation substantially alters the BM EC transcriptome compared to untreated controls, while CD45-targeted RIT severely remodels BM EC gene expression compared to non-targeted RIT.

**Figure 6:**
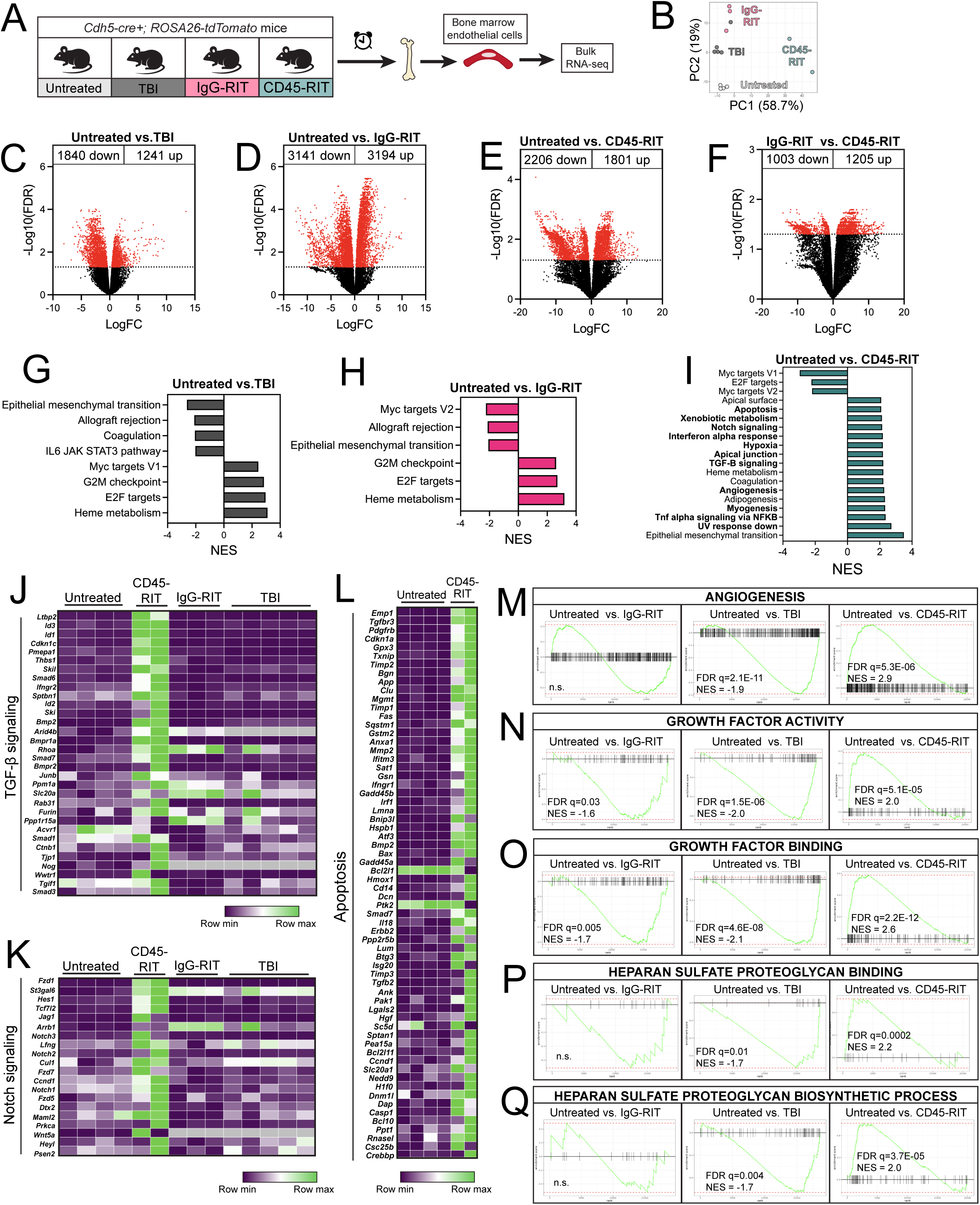
CD45-RIT transcriptionally reprograms bone marrow endothelial cells. **(A)** Study design depicting experimental pipeline to isolate BM ECs after treatment. **(B)** Principal component analysis scatter plot showing the samples from each group that were analyzed using bulk RNA sequencing. Volcano plot showing the fold change of differentially expressed genes when comparing untreated to **(C)** TBI, **(D)** IgG-RIT, or **(E)** CD45-RIT, or **(F)** IgG-RIT compared to CD45-RIT. Hallmarks identified with Gene Set Enrichment Analysis as being significantly different and greater than twofold enriched or suppressed in untreated samples compared to **(G)** TBI, **(H)** IgG-RIT, or **(I)** CD45-RIT. Heatmap showing differential expression of genes within the **(J)** TGF-β signaling, **(K)** Notch signaling, and **(L)** apoptosis hallmarks. Gene Set Enrichment Analysis was performed, and enrichment score plots from gene ontology terms for **(M)** angiogenesis, **(N)** growth factor activity, **(O)** growth factor binding, **(P)** heparan sulfate proteoglycan binding, and **(Q)** heparan sulfate proteoglycan biosynthesis are shown comparing untreated to TBI or untreated to CD45-RIT. (n=2-5 mice per condition from n=2 independent experiments, statistics show adjusted p-values).

Several gene sets were differentially enriched in ECs from TBI, IgG-RIT, or CD45-RIT-treated mice compared to untreated mice (**Figure 6G-I**). Notably, numerous gene sets, including those associated with apoptosis, Notch signaling, and TGFβ signaling, were significantly enriched in CD45-RIT ECs compared to untreated ECs but not in TBI or IgG-RIT compared to untreated ECs (**Figure 6I**, bolded terms). Heatmap analyses showing genes composing TGFβ and Notch signaling terms highlight specific gene expression changes in ECs after CD45-RIT that are not recapitulated upon IgG-RIT or TBI (**Figure 6J-K**). Genes associated with apoptosis were also dysregulated in ECs from CD45-RIT-treated mice compared to untreated (**Figure 6L**). These findings suggest that CD45-targeted RIT causes unique gene expression changes in BM ECs not seen in non-targeted RIT or TBI.

Using gene ontology, we analyzed how CD45-RIT impacted gene sets corresponding to molecular functions and biological processes. Compared to untreated ECs, gene sets associated with the regulation of angiogenesis were negatively enriched in ECs from TBI-treated mice and unchanged in IgG-RIT-treated mice. In contrast, gene sets associated with angiogenesis were positively enriched in CD45-RIT-treated compared to untreated ECs (**Fig. 6M**). Gene sets associated with growth factor activity and binding were negatively enriched in TBI or IgG-RIT-treated ECs compared to untreated, while they were positively enriched in CD45-RIT-treated ECs (**Fig. 6N-O**). Therefore, the radiation source and delivery method distinctly regulate the EC transcriptome.

Finally, as previous works identified critical roles for glycans, proteoglycans, and the glycocalyx in vascular functions [29–31], we assessed how CD45-targeted RIT affects gene sets pertaining to these processes. Gene ontology analysis revealed significant positive enrichment in gene sets pertaining to glycosaminoglycan binding, proteoglycan binding, heparan sulfate proteoglycan binding, and heparan sulfate biosynthetic process in ECs from CD45-RIT treated mice compared to controls. Genes that play key roles in heparan sulfate proteoglycan binding and heparan sulfate biosynthetic process were significantly negatively enriched in IgG-RIT ECs compared to untreated ECs, while proteoglycan and heparan sulfate proteoglycan binding were unchanged (**Figure 6P-Q**). These findings suggest that CD45-targeted RIT substantially remodels the transcriptional machinery curating the EC proteoglycome, which may be involved in controlling the regeneration of the vascular or hematopoietic system upon radiation injury.

## DISCUSSION

Our data indicate that CD45-targeted RIT using ^211^At depletes HSCs as effectively as TBI using ^137^Cs, but CD45- targeted RIT sustains low HSC levels until day +14 after treatment (**Figure 4**). Murine models showed that residual host HSCs negatively impact donor HSC engraftment upon transplant due to competition [32, 33]. Therefore, we postulate that CD45-targeted RIT as a pre-conditioning regimen before HCT may better support transplant success compared to non-targeted approaches. Consistent with this concept, non-human primate studies using CD45-targeted ^211^At RIT as a myeloablative preconditioning regimen were successful for HSPC gene therapy approaches, showcasing the promise of this technique [34]. Several outstanding questions remain, including whether ^211^At RIT and ^137^Cs TBI reduce HSC levels via similar cell-intrinsic or extrinsic molecular mechanisms, and whether CD45-targeted RIT causes secondary malignancies or organ damage [35, 36].

We show transient myelosuppression after radiation, which is prolonged with CD45 targeting compared to non-targeted IgG-RIT or TBI. One of the earliest alpha-emitter-based anti-CD45-RIT studies treated dogs with ^213^Bi- anti-canine CD45-RIT and reported no toxic effects. There was a mild, reversible blood count suppression where the timing of lymphocytic nadirs was more variable, whereas neutropenia and thrombocytopenia were more consistently most severe around 10 and approximately 12 days after RIT, respectively [37]. Subsequent dose-finding studies with the longer-lived alpha-emitter ^211^At used with anti-CD45-RIT dosed in dogs without HCT rescue also demonstrated dose-dependent myelosuppression and subsequent autologous recovery; the authors found that lymphopenia nadirs preceded neutropenic nadirs though there was more individual variability in lymphopenia than neutropenia [38]. When ^211^At was employed with anti-murine CD45-RIT in immunocompetent mice, leukocytes were most affected at about two weeks post-treatment but recovered by four. However, the prior study did not pursue neutropenic and lymphopenic kinetics with the same granularity as ours [14].

Another critical question is whether anti-CD45-RIT preferentially targets one leukocyte compartment over others and its potential for lymphodepletion, for example, prior to CAR T-cell therapy. We show that lymphopenia manifests as early as 24 hours after RIT, whereas the neutropenia nadir occurs between days +4-7, as seen in prior anti-CD45-RIT studies in canine [7, 39] and murine models [40]. Our BM analyses show rapidly decreased BM cellularity by day +1 after anti-CD45-RIT, persisting through day 14 compared to TBI or IgG-RIT (**Figure 3**). The CD3^+^ marrow compartment is maximally depressed by day +1 after RIT, whereas the Mac-1/Gr-1^+^ nadir is closer to day +4-7 before recovering between days +14 and +28. To what extent the tempo of nadirs can be modulated by radioactivity dose or BM target is yet to be explored, as these studies only used a standard, tolerated dose of 20µCi/mouse.

The primary benefit of RIT compared to non-targeted approaches is thought to occur from improved radiation delivery to target cells [41, 42]. Our data imply the BM niche also responds differently to targeted RIT compared to non-targeted radiation. For example, RNA sequencing revealed that BM ECs are enriched for the interferon alpha (IFN-α) signal response upon RIT (**Figure 6**). IFN-α therapy has had promising results for leukemia patients; IFN-α promotes cell killing directly or indirectly by supporting anticancer immunomodulation [43–46]. We postulate that RIT-mediated activation of IFN-α signaling in the BM niche could enhance leukemia killing by supporting both radiation- and immunomodulatory-mediated killing. Future studies will test this possibility directly by dissecting the impact of RIT on immune cell recruitment to the BM for cancer cell clearance.

Prior works showed the BM niche is restructured by TBI [22, 25, 27]. Our work highlights that the BM niche is architecturally remodeled by CD45-targeted RIT with eventual restoration of vascular order (**Figure 5**). We suggest that this may occur either due to the close physical proximity of ECs to CD45-expressing cells or indirect effects incurred by ECs when their microenvironment is transformed by hematologic depletion. RNA sequence analysis revealed gene sets enriched in ECs upon CD45-RIT treatment compared to untreated conditions, such as TGFβ and Notch signaling; gene sets associated with these pathways were not enriched in ECs from TBI or IgG-RIT-treated mice. Published reports showcase the importance of TGFβ and Notch signaling in vascular and hematologic regeneration after injury [23, 47–49]. Whether EC activation of these transcriptional pathways is beneficial to hematopoietic and vascular regeneration after RIT is unknown. Further, ECs may activate these pathways to restrain regeneration, for example, to ensure DNA damage is repaired before differentiation occurs. Future work also needs to clarify the impact of CD45-targeting RIT on other critical components of the BM niche known to be remodeled by stress, like mesenchymal stromal cells and adipocytes [22, 50].

This detailed description of PB and BM responses to ^211^At-anti-CD45-RIT in immunocompetent mouse models and how it compares to non-targeted therapeutic approaches is timely, given the recent growth in radiopharmaceuticals, especially in treating hematologic malignancies [51–53] . With efficacy reports of ^211^At- based RIT in preclinical animal models of non-Hodgkin lymphoma, AML, and multiple myeloma, [54–56] a first in human ^211^At-anti-CD45-RIT prior to allogeneic HCT in high-risk myeloid malignancies trial was initiated. Preliminary results of the first 20 patients treated on the dose escalation portion showed an estimated 1-year overall survival of 43%, [8] which is impressive considering that 16 patients had measurable residual disease or >5% blasts in the BM where HCT is known to be less effective [57–59]. Our study highlights the need to define the short- and long-term tissue, cellular, and molecular consequences to the human BM vasculature, as these remodeling events may influence patient outcomes.

## ACKNOWLEDGMENTS

We are grateful to Cyd McKay and Christina Root for their assistance with animal studies and Donald Hamlin and Yawen Li for synthesizing and facilitating the procurement of the radiolabeled antibodies used in this report. The production of astatine-211 at the University of Washington was supported by the U.S. Department of Energy’s Isotope Program, managed by the Office of Science for Isotope R&D and Production. We would like to acknowledge the excellent assistance provided by the Fred Hutchinson Cancer Center Shared Resources, such as the Comparative Medicine, Experimental Histopathology (Audrey Heintz and Stephanie Weaver), Flow Cytometry, Antibody Production, and Genomics & Bioinformatics Shared Resources (Feinan Wu). We thank Julia Sober, Shawnie Zahniser, and Susan Parazzoli for their assistance with radiation safety-related affairs. The investigation was supported in part by parent grant and administrative supplement 3R21CA283589 to JJO and CMT and 5K01DK126989 to CMT from the National Institutes of Health, USA. CMT is also supported by 23CDA1039196 from the American Heart Association and a New Investigator Award from the American Society for Transplantation and Cellular Therapy. KAW was supported by a Fellowship from the Fred Hutchinson Cancer Center Office of Faculty Affairs and the Graduate Research Fellowship Program from the National Science Foundation. This research was supported by NIH P30 CA015704 to the Fred Hutch/University of Washington/Seattle Children’s Cancer Consortium, which includes the Comparative Medicine, Experimental Histopathology, Flow Cytometry, Antibody Production, and Genomics & Bioinformatics Shared Resources.

## AUTHORSHIP CONTRIBUTIONS

Project conception (CMT and JJO) performed experiments (MWH, NJS, KAW, SD, FG), analyzed data (MWH, CMT), made figures (CMT and MWH), wrote the manuscript (CMT, MWH, JJO).

## CONFLICT OF INTEREST DISCLOSURES

The authors declare no competing financial interests.

## Notes

https://www.ncbi.nlm.nih.gov/geo/query/acc.cgi?acc=GSE293137

### Competing Interest Statement

The authors have declared no competing interest.

## REFERENCES

1. Dee, E.C., et al., Radiotherapy for haematological malignancies. Lancet Haematol, 2024. 11(10): p. e721–e722.

2. Mikhaeel, N.G., et al., The Optimal Use of Imaging in Radiation Therapy for Lymphoma: Guidelines from the International Lymphoma Radiation Oncology Group (ILROG). Int J Radiat Oncol Biol Phys, 2019. 104(3): p. 501–512.

3. Junghanss, C., et al., Incidence and outcome of bacterial and fungal infections following nonmyeloablative compared with myeloablative allogeneic hematopoietic stem cell transplantation: a matched control study. Biol Blood Marrow Transplant, 2002. 8(9): p. 512–20.

4. Ozdemir, Z.N. and S. Civriz Bozdag, Graft failure after allogeneic hematopoietic stem cell transplantation. Transfus Apher Sci, 2018. 57(2): p. 163–167.

5. Feinendegen, L.E. and J.J. McClure, *Alpha-emitters for medical therapy: workshop of the United States Department of Energy: Denver, Colorado, May 30-31*, *1996*. Radiation Research, 1997. 148(2): p. 195-201.

6. Humm, J.L., A microdosimetric model of astatine-211 labeled antibodies for radioimmunotherapy. Int J Radiat Oncol Biol Phys, 1987. 13(11): p. 1767–73.

7. Frost, S.H.L., et al., (211)At-Labeled Anti-CD45 Antibody as a Nonmyeloablative Conditioning for Canine DLA-Haploidentical Stem Cell Transplantation. J Nucl Med, 2024. 65(9): p. 1443–1449.

8. Sandmaier, B.M., et al., 57 - A Phase I Trial of First-in-Human Alpha-Emitter Astatine-211-Labeled Anti- CD45 Antibody (211At-BC8-B10) in Combination with Fludarabine and TBI As Conditioning for Allogeneic Hematopoietic Cell Transplantation (HCT) for Patients with Refractory/Relapsed Leukemia or High-Risk Myelodysplastic Syndrome (MDS): Preliminary Results of Dose Escalation. Transplantation and Cellular Therapy, 2021. 27(3, Supplement): p. S54.

9. *A Study Evaluating Escalating Doses of 211^At-Labeled Anti-CD45 MAb BC8-B10 (211^At-BC8-B10) Followed by Allogeneic Hematopoietic Cell Transplantation for High-Risk Acute Myeloid Leukemia (AML), Acute Lymphoblastic Leukemia (ALL), or Myelodysplastic Syndrome (MDS)*, I. National Cancer, Editor. 2017.

10. Albertsson, P., et al., Astatine-211 based radionuclide therapy: Current clinical trial landscape. Front Med (Lausanne), 2022. 9: p. 1076210.

11. *A Phase I/II Study Evaluating Escalating Doses of 211At-Labeled Anti-CD45 MAb BC8-B10 (211At- BC8-B10) Followed by Related Haplo-Identical Allogeneic Hematopoietic Cell Transplantation for High-Risk Acute Leukemia or Myelodysplastic Syndrome (MDS)*, I. National Cancer, Editor. 2018.

12. *A Phase I Trial Evaluating Escalating Doses of ²¹¹At-Labeled Anti-CD38 Monoclonal Antibody Followed by HLA-Matched or Haploidentical Donor Hematopoietic Cell Transplantation for High-Risk Multiple Myeloma*, I. National Cancer, Editor. 2020.

13. Wilbur, D.S., et al., Biotin reagents for antibody pretargeting. 5. Additional studies of biotin conjugate design to provide biotinidase stability. Bioconjug Chem, 2001. 12(4): p. 616-23.

14. Orozco, J.J., et al., Anti-CD45 radioimmunotherapy using (211)At with bone marrow transplantation prolongs survival in a disseminated murine leukemia model. Blood, 2013. 121(18): p. 3759–67.

15. Amend, S.R., K.C. Valkenburg, and K.J. Pienta, Murine Hind Limb Long Bone Dissection and Bone Marrow Isolation. J Vis Exp, 2016(110).

16. Himburg, H.A., et al., Distinct Bone Marrow Sources of Pleiotrophin Control Hematopoietic Stem Cell Maintenance and Regeneration. Cell Stem Cell, 2018. 23(3): p. 370–381 e5.

17. Robinson, M.D. and A. Oshlack, A scaling normalization method for differential expression analysis of RNA-seq data. Genome Biol, 2010. 11(3): p. R25.

18. Subramanian, A., et al., Gene set enrichment analysis: a knowledge-based approach for interpreting genome-wide expression profiles. Proc Natl Acad Sci U S A, 2005. 102(43): p. 15545–50.

19. Domen, J., S.H. Cheshier, and I.L. Weissman, The role of apoptosis in the regulation of hematopoietic stem cells: Overexpression of Bcl-2 increases both their number and repopulation potential. J Exp Med, 2000. 191(2): p. 253–64.

20. Domen, J., K.L. Gandy, and I.L. Weissman, Systemic overexpression of BCL-2 in the hematopoietic system protects transgenic mice from the consequences of lethal irradiation. Blood, 1998. 91(7): p. 2272–82.

21. Osawa, M., et al., Long-term lymphohematopoietic reconstitution by a single CD34-low/negative hematopoietic stem cell. Science, 1996. 273(5272): p. 242-5.

22. Fang, S., et al., VEGF-C protects the integrity of the bone marrow perivascular niche in mice. Blood, 2020. 136(16): p. 1871–1883.

23. Guo, P., et al., Endothelial jagged-2 sustains hematopoietic stem and progenitor reconstitution after myelosuppression. J Clin Invest, 2017. 127(12): p. 4242–4256.

24. Hooper, A.T., et al., Engraftment and reconstitution of hematopoiesis is dependent on VEGFR2- mediated regeneration of sinusoidal endothelial cells. Cell Stem Cell, 2009. 4(3): p. 263–74.

25. Chen, Q., et al., Apelin(+) Endothelial Niche Cells Control Hematopoiesis and Mediate Vascular Regeneration after Myeloablative Injury. Cell Stem Cell, 2019. 25(6): p. 768–783 e6.

26. Butler, J.M., et al., Endothelial cells are essential for the self-renewal and repopulation of Notch-dependent hematopoietic stem cells. Cell stem cell, 2010. 6(3): p. 251–264.

27. Termini, C.M., et al., Neuropilin 1 regulates bone marrow vascular regeneration and hematopoietic reconstitution. Nature Communications, 2021. 12(1).

28. Himburg, H.A., et al., A Molecular Profile of the Endothelial Cell Response to Ionizing Radiation. Radiat Res, 2016. 186(2): p. 141–52.

29. Corti, F., et al., Syndecan-2 selectively regulates VEGF-induced vascular permeability. Nat Cardiovasc Res, 2022. 1(5): p. 518–528.

30. Corti, F., et al., N-terminal syndecan-2 domain selectively enhances 6-O heparan sulfate chains sulfation and promotes VEGFA(165)-dependent neovascularization. Nat Commun, 2019. 10(1): p. 1562.

31. Queisser, K.A., et al., COVID-19 generates hyaluronan fragments that directly induce endothelial barrier dysfunction. JCI Insight, 2021. 6(17).

32. Bhattacharya, D., et al., Niche recycling through division-independent egress of hematopoietic stem cells. J Exp Med, 2009. 206(12): p. 2837–50.

33. Czechowicz, A., et al., Efficient transplantation via antibody-based clearance of hematopoietic stem cell niches. Science, 2007. 318(5854): p. 1296-9.

34. Radtke, S., et al., CD45-Directed Radioimmunotherapy with the Alpha-Emitter Astatine-211 As Conditioning for Autologous Hematopoietic Stem/Progenitor Cell Gene Therapy: Results from a Nonhuman Primate Model. Blood, 2024. 144: p. 2014.

35. Bearman, S.I., et al., Regimen-related toxicity in patients undergoing bone marrow transplantation. J Clin Oncol, 1988. 6(10): p. 1562–8.

36. Matthes-Martin, S., et al., Organ toxicity and quality of life after allogeneic bone marrow transplantation in pediatric patients: a single centre retrospective analysis. Bone Marrow Transplant, 1999. 23(10): p. 1049–53.

37. Sandmaier, B.M., et al., Bismuth 213-labeled anti-CD45 radioimmunoconjugate to condition dogs for nonmyeloablative allogeneic marrow grafts. Blood, 2002. 100(1): p. 318–26.

38. Chen, Y., et al., Durable donor engraftment after radioimmunotherapy using alpha-emitter astatine-211- labeled anti-CD45 antibody for conditioning in allogeneic hematopoietic cell transplantation. Blood, 2012. 119(5): p. 1130–8.

39. Frost, S.H., et al., *alpha-Imaging Confirmed Efficient Targeting of CD45-Positive Cells After 211At- Radioimmunotherapy for Hematopoietic Cell Transplantation*. J Nucl Med, 2015. 56(11): p. 1766–73.

40. Nakamae, H., et al., Biodistributions, myelosuppression, and toxicities in mice treated with an anti- CD45 antibody labeled with the alpha-emitting radionuclides bismuth-213 or astatine-211. Cancer Res, 2009. 69(6): p. 2408–15.

41. Walter, R.B., Where do we stand with radioimmunotherapy for acute myeloid leukemia? Expert Opin Biol Ther, 2022. 22(5): p. 555–561.

42. Ivanov, A., R. Swann, and T. Illidge, New insights into the mechanisms of action of radioimmunotherapy in lymphoma. J Pharm Pharmacol, 2008. 60(8): p. 987–98.

43. Mo, X.D., et al., Interferon-alpha Is Effective for Treatment of Minimal Residual Disease in Patients with t(8;21) Acute Myeloid Leukemia After Allogeneic Hematopoietic Stem Cell Transplantation: Results of a Prospective Registry Study. Oncologist, 2018. 23(11): p. 1349–1357.

44. Mo, X.D., et al., IFN-alpha Is Effective for Treatment of Minimal Residual Disease in Patients with Acute Leukemia after Allogeneic Hematopoietic Stem Cell Transplantation: Results of a Registry Study. Biol Blood Marrow Transplant, 2017. 23(8): p. 1303–1310.

45. Jiang, H., et al., Interferon-alpha as maintenance therapy can significantly reduce relapse in patients with favorable-risk acute myeloid leukemia. Leuk Lymphoma, 2021. 62(12): p. 2949–2956.

46. Kiladjian, J.J., R.A. Mesa, and R. Hoffman, The renaissance of interferon therapy for the treatment of myeloid malignancies. Blood, 2011. 117(18): p. 4706–15.

47. Zhao, G., et al., TGF-betaR2 signaling coordinates pulmonary vascular repair after viral injury in mice and human tissue. Sci Transl Med, 2024. 16(732): p. eadg6229.

48. Brenet, F., et al., TGFbeta restores hematopoietic homeostasis after myelosuppressive chemotherapy. J Exp Med, 2013. 210(3): p. 623–39.

49. Shao, L., et al., A Tie2-Notch1 signaling axis regulates regeneration of the endothelial bone marrow niche. Haematologica, 2019. 104(11): p. 2164–2177.

50. Zhou, B.O., et al., Bone marrow adipocytes promote the regeneration of stem cells and haematopoiesis by secreting SCF. Nat Cell Biol, 2017. 19(8): p. 891–903.

51. Mawad, R., et al., Radiolabeled anti-CD45 antibody with reduced-intensity conditioning and allogeneic transplantation for younger patients with advanced acute myeloid leukemia or myelodysplastic syndrome. Biol Blood Marrow Transplant, 2014. 20(9): p. 1363–8.

52. Gyurkocza, B., et al., Randomized Phase III SIERRA Trial of (131)I-Apamistamab Before Allogeneic Hematopoietic Cell Transplantation Versus Conventional Care for Relapsed/Refractory AML. J Clin Oncol, 2025. 43(2): p. 201–213.

53. Pagel, J.M., et al., Allogeneic hematopoietic cell transplantation after conditioning with 131I-anti-CD45 antibody plus fludarabine and low-dose total body irradiation for elderly patients with advanced acute myeloid leukemia or high-risk myelodysplastic syndrome. Blood, 2009. 114(27): p. 5444–53.

54. Frost, S., et al., Anti-CD45 monoclonal antibody (MAb) dose optimization for astatine-211 ^211^At)- radioimmunotherapy (RIT) of relapsed non-Hodgkin lymphoma (NHL) in a canine model. Journal of Nuclear Medicine, 2014. 55(supplement 1): p. 637–637.

55. Laszlo, G.S., et al., Development of [(211)At]astatine-based anti-CD123 radioimmunotherapy for acute leukemias and other CD123+ malignancies. Leukemia, 2022. 36(6): p. 1485–1491.

56. O’Steen, S., et al., The alpha-emitter astatine-211 targeted to CD38 can eradicate multiple myeloma in a disseminated disease model. Blood, 2019. 134(15): p. 1247–1256.

57. Ali, N., et al., Measurable residual disease as predictor of post-day +100 relapses after allografting in adult AML. Blood Adv, 2025. 9(3): p. 558–570.

58. Gang, M., M. Othus, and R.B. Walter, Significance of Measurable Residual Disease in Patients Undergoing Allogeneic Hematopoietic Cell Transplantation for Acute Myeloid Leukemia. Cells, 2025. 14(4).

59. Orvain, C., et al., Relationship between donor source, pre-transplant measurable residual disease, and outcome after allografting for adults with acute myeloid leukemia. Leukemia, 2025. 39(2): p. 381–390.

