## Supplemental Table 1 for "The bone marrow niche and hematopoietic system are distinctly remodeled by CD45-targeted astatine-211 radioimmunotherapy"

**Supplementary Table 1: Manuscript Resources table.** Vendor information for antibodies, assays, mouse models, and software used throughout manuscript.

| ANTIBODIES | | | | |
|---|---|---|---|---|
| **Name** | **Vendor** | **Catalog Number** | **Clone** | **Dilution** |
| V450 Mouse Lineage Antibody Cocktail with isotype | BD Biosciences | 561301 | n/a | 20µL / $10^6$ cells |
| APC/Cyanine 7 anti-mouse Ly-6A/E | Biolegend | 108125 | D7 | 1µg / $10^6$ cells |
| APC/Cyanine 7 Rat IgG2a, κ Isotype control | Biolegend | 400523 | RTK2758 | 1µg / $10^6$ cells |
| PE anti-mouse CD117 | Biolegend | 105807 | 2B8 | 1µg / $10^6$ cells |
| PE Rat IgG2b, κ Isotype control | Biolegend | 400607 | RTK4530 | 1µg / $10^6$ cells |
| PE/Cyanine 7 anti-mouse CD34 | Biolegend | 119325 | MEC14.7 | 1µg / $10^6$ cells |
| PE/Cyanine 7 Rat IgG2a, κ Isotype control | Biolegend | 400521 | RTK2758 | 1µg / $10^6$ cells |
| KIRAVIA Blue 520 anti-mouse CD135 | Biolegend | 135323 | A2F10 | 1µg / $10^6$ cells |
| KIRAVIA Blue 50 Rat IgG2a, κ isotype control | Biolegend | 400575 | RTK2758 | 1µg / $10^6$ cells |
| Brilliant Violet 605 anti-mouse CD150 | Biolegend | 115927 | TC15-12F12.2 | 1µg / $10^6$ cells |
| Brilliant Violet 605 Rat IgG2a, κ isotype control | Biolegend | 400539 | RTK2758 | 1µg / $10^6$ cells |
| Alexa Fluor 700 anti-mouse CD48 | Biolegend | 103425 | HM48-1 | 1µg / $10^6$ cells |
| Hamster IgG Isotype Ctrl | Biolegend | 400926 | HTK888 | 1µg / $10^6$ cells |
| APC Rat Anti-Mouse TER-119 / Erythroid cells | BD Biosciences | 557909 | TER-119 | 0.4µg / $10^5$ cells |
| APC Rat IgG2b, κ isotype control | BD Biosciences | 553991 | | 1µg / $10^6$ cells |
| Brilliant Violet 421 anti-mouse CD3 | Biolegend | 100228 | 17A2 | 0.4µg / $10^5$ cells |
| Brilliant Violet 421 Rat IgG2b, κ Isotype Control | Biolegend | 400639 | RTK4530 | 1µg / $10^6$ cells |
| BV711 Rat Anti-Mouse CD45R/B220 | BD Biosciences | 563892 | RA3-6B2 | 0.4µg / $10^5$ cells |
| BV711 Hamster IgG1, κ isotype control | BD Biosciences | 563128 | | 1µg / $10^6$ cells |
| PE Rat Anti-CD11b | BD Biosciences | 553311 | M1/70 | 0.4µg / $10^5$ cells |
| PE Rat Anti-Mouse Ly-6G and Ly-6C | BD Biosciences | 553128 | RB6-6C5 | 0.4µg / $10^5$ cells |
| PE Rat IgG2b, κ Isotype control | BD Biosciences | 553989 | | 1µg / $10^6$ cells |
| Purified Rat Anti-Mouse CD16/CD32 (Mouse BD FC Block) | BD Biosciences | 553142 | 2.4G2 | 2.5µg / $10^6$ cells |
| Alexa Fluor 700 Rat Anti-Mouse CD45 | Biolegend | 103128 | 30-F11 | 1µg / $10^6$ cells |
| Alexa Fluor 700 Rat IgG2b, κ isotype control | Biolegend | 400628 | RTK4530 | 1µg / $10^6$ cells |
| CD31 | Cell signaling technologies | 77699S | | 1:200 |
| Rat IgG2b isotype control | BioXCell | BE0090 | LTF-2 | |
| **MOUSE MODELS** | | | | |

| Name | Vendor | Catalog Number |
|---|---|---|
| Mouse C57BL/6J | The Jackson Laboratory | 000664 |
| Mouse CDH5 Cre (B6.FVB-Tg(Cdh5-cre)7Mlia/J) | The Jackson Laboratory | 6137 |
| Mouse TdTomato (B6.Cg-Gt(ROSA)26Sortm14(CAG-tdTomato)Hze/J) | The Jackson Laboratory | 7914 |
| **REAGENTS** | | |
| Iscove's Modified Dulbecco's Medium | Gibco | 12440-053 |
| Penicillin-Streptomycin Solution | Fisher Scientific | MT30002CI |
| Fetal bovine serum | Fisher Scientific | 501527078 |
| Compensation beads | Fisher Scientific | 3031-5813-28 |
| 7AAD cell viability stain | BD Biosciences | 559925 |
| Methocult GF M3434 | Stemcell Technologies | 03434 |
| SmartDish | Stemcell Technologies | 27371 |
| EDTA vacutainers | BD | 101005 |
| Ammonium-Chloride-Potassium (ACK) Lysing Buffer | Fisher Scientific | 50-983-219 |
| Liberase | Millipore Sigma | 5401020001 |
| **COMMERCIAL KITS** | | |
| **Name** | **Vendor** | **Catalog Number** |
| Direct lineage cell depletion kit mouse | Miltenyi | 130-110-470 |
| LS columns | Miltenyi | 130-042-401 |
| RNeasy Micro Kit | Qiagen | 74004 |
| **SOFTWARE** | | |
| **Name** | **Vendor** | **Catalog Number** |
| HALO v.3.6.4134.265 | Indica Labs | |
| FlowJo v10 | BD Biosciences | n/a |
| GraphPad Prism Version 10.4.1 (532) | GraphPad Software | |
| STEMVision Analyzer v2.6.7.0 | Stemcell Technologies | |
| STEMVision Colony Marker v2.5.0.0 | Stemcell Technologies | |
| R v4.4.2 | R foundation for statistical computing | |
| STAR v2.7.7a | Dobin et al. 2013 | Dobin A, Davis CA, Schlesinger F, et al. STAR: ultrafast universal RNA-seq aligner. Bioinformatics. 2013;29(1):15-21. doi:10.1093/bioinformatics/bts635 |
| FastQC v0.11.9 | Babraham Bioinformatics | |
| RNA-SeQC v2.3.4 | DeLuca et al. 2012 | DeLuca DS, Levin JZ, Sivachenko A, et al. RNA-SeQC: RNA-seq metrics for quality control and process optimization. Bioinformatics. 2012;28(11):1530-1532. doi:10.1093/bioinformatics/bts196 |

| | | |
|---|---|---|
| RSeQC v4.0.0 | Wang et al. 2012 | Wang L, Wang S, Li W. RSeQC: quality control of RNA-seq experiments. Bioinformatics. 2012;28(16):2184-2185. doi:10.1093/bioinformatics/bts356 |
| Bioconductor edgeR v3.36.0 | Robinson et al. 2010 | Robinson MD, McCarthy DJ, Smyth GK. edgeR: a Bioconductor package for differential expression analysis of digital gene expression data. Bioinformatics. 2010;26(1):139-140. doi:10.1093/bioinformatics/btp616 |
| Fgea | Korotkevich et al. 2021 | Fast gene set enrichment analysis Gennady Korotkevich, Vladimir Sukhov, Nikolay Budin, Boris Shpak, Maxim N. Artyomov, Alexey Sergushichev bioRxiv 060012; doi: https://doi.org/10.1101/060012 |
